# Unbiased *in vivo* exploration of nuclear bodies-enhanced sumoylation reveals that PML orchestrates embryonic stem cell fate

**DOI:** 10.1101/2021.06.29.450368

**Authors:** Sarah Tessier, Omar Ferhi, Marie-Claude Geoffroy, Roman Gonzalez-Prieto, Antoine Canat, Samuel Quentin, Marika Pla, Michiko Niwa-Kawakita, Pierre Bercier, Domitille Rérolle, Pierre Therizols, Emmanuelle Fabre, Alfred C.O. Vertegaal, Hugues de Thé, Valérie Lallemand-Breitenbach

## Abstract

Membrane-less organelles are condensates formed by phase separation whose functions often remain enigmatic. Upon oxidative stress, PML scaffolds Nuclear Bodies (NBs) to regulate senescence or metabolic adaptation, but their role in pluripotency remains elusive. Here we establish that PML is required for basal SUMO2/3 conjugation in mESCs and oxidative stress-driven sumoylation in mESCs or *in vivo*. PML NBs create an oxidation-protective environment for UBC9-driven SUMO2/3 conjugation of PML partners, often followed by their poly-ubiquitination and degradation. Differential *in vivo* proteomics identified several members of the KAP1 complex as PML NB-dependent SUMO2-targets. The latter drives functional activation of this key epigenetic repressor. Accordingly, *Pml^−/−^* mESCs re-express transposable elements and display features of totipotent-like cells, a process further enforced by PML-controlled SUMO2-conjugation of DPPA2. Finally, PML is required for adaptive stress responses in mESCs. Collectively, PML orchestrates mESC fate through SUMO2-conjugation of key transcriptional or epigenetic regulators, raising new mechanistic hypotheses about PML roles in normal or cancer stem cells.

## Introduction

Membrane-less organelles are stress-sensitive deposits of concentrated bio-molecules. One specific type of nuclear body (NB) is scaffolded by the promyelocytic leukemia protein (PML) and is required for essential processes such as senescence, metabolism or viral restriction^1,2^. Restoration of PML NBs with arsenic or retinoic acid therapies underlies cures of patients with acute promyelocytic leukemia (APL), emphasizing their physio-pathological relevance^3^. *In vivo*, PML senses reactive oxygen species (ROS), driving PML NB assembly and physiological responses to oxidative stress^4^. Mimicking ROS, arsenic directly binds PML, promoting its multimerization and NB assembly^5–8^. PML is also essential for the fitness of normal or malignant stem cells^9–12^ and PML down-regulation alleviates reprograming of mouse embryonic fibroblast (MEFs) into iPSCs^13^. These findings raise the question of how PML NBs contribute to stem cell biology and their response to stress.

The sumoylated PML scaffold attracts many disparate partner proteins inside NBs. However, PML NB insolubility has thwarted biochemical purification efforts to establish a comprehensive list of these partners and no general biochemical activity has been demonstrated so far that might explain the diversity of PML-regulated processes^1,2^. PML NBs have a proposed role in post-translational modifications of partners, in particular sumoylation, since UBC9, the SUMO-E2 enzyme, concentrates within NBs^1,2,8^. Functionally, conjugation by SUMO2/3 (two indistinguishable SUMO paralogues) may control proteasomal degradation through recruitment of SUMO-targeted ubiquitin ligases^6,14–17^. Sumoylation may also regulate transcription^18^ and sustain epigenetic modifications, notably those essential for maintaining the identity of mouse embryonic stem cells (mESCs)^19–21^. In various cell lines, oxidative stress increases sumoylation, while paradoxically SUMO enzymes are inhibited by ROS^22–24^. These observations raise the question of whether SUMO might be involved in some PML-mediated functions, in particular in stem cell fate.

Here we establish that arsenic-induced PML NB biogenesis drives sumoylation and we identify the first unbiased target list. Remarkably, in mESCs, PML favors SUMO2 conjugation of the KAP1 epigenetic regulatory complex, as well as that of the master transcription factor DPPA2 to oppose 2-Cell-Like (2CL) state, unraveling an unexpected key role for PML in the homeostasis of pluripotent ESCs.

## Results

### PML mediates stress-induced mESC growth inhibition and SUMO2 conjugation *in vivo*

In pluripotent mESCs, PML expression is high and NBs are abundant compared to their committed state upon retinoic acid treatment and LIF withdraw (Fig. 1a). Arsenic treatment rapidly increased PML NB assembly and recruited SUMO1/2/3 (Supplementary Fig. 1a), as in transformed cell lines^5^. To functionally explore any role of PML NBs in mESC homeostasis and stress response, we generated CRISPR/Cas9-engineered *Pml* knock out (*Pml^−/−^*) mESCs, and assessed arsenic response in two independent clones. Critically, PML was required for arsenic-driven growth arrest (Fig. 1b), implying that PML dynamics plays a key role in stress response of mESCs.

**Figure 1.**
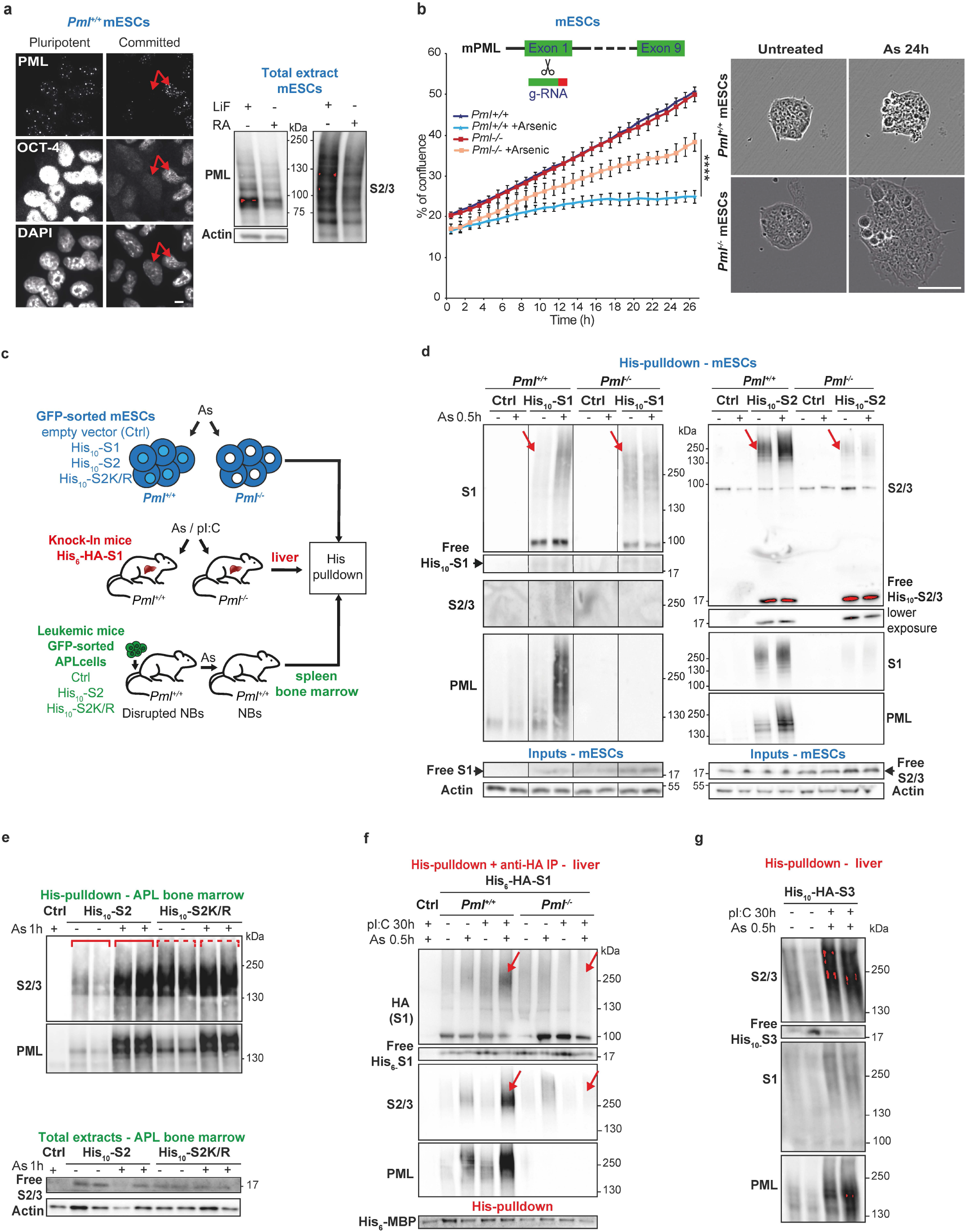
PML is required for stress response and sumoylation in mESCs and mice. **(a)** PML NBs (representative confocal analysis, left) expression (Western blot analysis, right) and in pluripotent (LIF) versus committed (Retinoic Acid and LIF withdrawal) mESCs, in which Oct4 decreases (compare arrowed cell). Scale: 5um. **(b)** IncuCyte cell proliferation assay showing arsenic stress-resistance with loss of Pml, with schematic of Pml edition. Mean values of n=3 +/− s.d., ****p<0.0001, Mann-Whitney test (left), and representative images (right), Scale: 50μm. Representative of 2 CrispR/Cas9-generated *Pml*^−/−^ mESC clones. **(c)** Experimental mouse models with tunable PML NBs: (Blue) *Pml*^+/+^ or *Pml*^−/−^ mESCs stably expressing His_10_-SUMO1-, His_10_-SUMO2-, His_10_-SUMO2K/R-IRES-GFP or GFP (His_10_-S1, His_10_-S2, His_10_-S2K/R, Ctrl). (Red) *His_6_-HA-Sumo1* knock-in;*Pml*^+/+^ or *Pml*^−/−^ mice injected with arsenic and pI:C to maximize NBs. (Green) APL mice obtained from serial transplantations of APL cells (Ctrl APL) or sorted APL cells expressing the indicated His_10_-SUMOs, with arsenic injection to restore PML NBs. **(d)** Pulldown (PD) of His_10_-S1 (left) or His_10_-S2 (right) conjugates showing arsenic-increased sumoylation in *Pml*^+/+^ mESCs only. Inputs indicating similar low levels of free His_10_-SUMO expression. Arrows: baseline sumoylation depending on PML. Representative data, n>3. Supplementary Fig. 1e for statistics. **(e)** PD from APL bone marrows as in (1d), showing increases in His_10_-S2 but not His_10_-S2K/R conjugation upon arsenic injection. Total extracts indicate respective levels of free S2/3. Representative experiment (2 mice per condition, 1 Ctrl APL with untagged SUMO2), n=3. **(f)** His_6_-HA-S1 conjugates dually purified from liver (top), showing PML dependent increase in S1/2/3 conjugation upon arsenic/pI:C (red arrows). Ctrl: Pml+/+mice, His_6_-MBP recombinant protein: internal PD control. See Supplementary Fig. 1g for statistics. **(g)** Same as (1f), using His_10_-HA-S3 knockin mice.

Similar to PML expression, global SUMO conjugation was higher in mESCs than in committed cells (Fig. 1a, right). To explore any role of PML NBs in sumoylation control, we leveraged mESCs and two *in vivo* biological systems with tunable NB biogenesis by arsenic treatment. First, in *Pml^+/+^ and Pml^−/−^* mESCs, we stably expressed His_10_-SUMO1 or 2 (ref^25^) at low levels compared to endogenous SUMO peptides (Fig. 1c, d-inputs). Second, we expressed low His_10_-SUMO2 level in an APL mouse model, where PML is expressed, but NBs are disrupted, a phenotype rapidly reversed by arsenic^3,5^ (Fig.1c, Supplementary Fig. 1b). Third, using His_6_-HA-tagged *Sumo1* knock-in mice^26^, we derived *His_6_-HA-Sumo1;Pml^+/+^* and *His_6_-HA-Sumo1;Pml^−/−^* mice. We studied livers of these mice where PML expression can be boosted by polyI:C (pI:C)^27^ (Fig. 1c, Supplementary Fig. 1c) and arsenic further drives PML NBs biogenesis (Supplementary Fig. 1d). To assess basal and arsenic-induced sumoylation, we performed His-pulldown enrichment. First, in mESCs, arsenic rapidly promoted conjugation by SUMO1 and SUMO2/3 only in *Pml^+/+^* cells (Fig. 1d left and right respectively, quantification Supplementary Fig. 1e). Moreover, in untreated *Pml^−/−^* mESCs, basal His_10_-SUMO2 conjugation was less efficient than in *Pml^+/+^* mESCs and this baseline defect was compensated by enhanced SUMO1 conjugation (Fig. 1d, right panel arrowed). Similarly, in His_10_-SUMO2-APLs, we observed a drastic global SUMO2 hyper-conjugation (and to a lesser extend SUMO1) upon arsenic treatment (Fig. 1e, and Supplementary Fig. 1f), following NB restoration (Supplementary Fig. 1b). Finally, in the liver of *His_6_-HA-Sumo1* knock-in mice, arsenic treatment increased His_6_-HA-SUMO1 conjugation and also massively induced their SUMO2/3-modification, an effect sharply enhanced by pI:C priming (Fig. 1f, quantification Supplementary Fig. 1g). Here again, arsenic-enhanced sumoylation was not observed in the *Pml^−/−^* background. Preliminary experiments in *HA-His_10_-Sumo3* knock-in mice (encoding the SUMO2 paralogue) demonstrated that arsenic-enhancement of SUMO2 conjugation was of much greater amplitude than that of SUMO1 (Fig. 1g). Collectively, all three models converge to demonstrate that basal and stress-induced PML NB assembly correlates with global hyper-SUMO2 conjugation *in vivo*.

SUMO2/3 being prone to its own sumoylation^19^, we questioned PML contribution to the formation of SUMO2-containing chains upon stress. In both mESCs and APL mice, stable expression of a His_10_-SUMO2 mutant defective for chain formation (SUMO2K/R)^25^ blocked arsenic-increased global SUMO2 conjugation (Fig. 1e, Supplementary Fig. 1h), while arsenic-induced PML multi-sumoylation was preserved. Basal levels of global SUMO2 conjugates were sharply increased by His_10_-SUMO2K/R expression, likely reflecting significant baseline SUMO2 chain-mediated catabolism (Fig. 1e, right part). Collectively, these findings support the conclusion that PML drives basal and stress-induced sumoylation of protein partners, primarily by SUMO2/3 *in vivo*.

### UBC9 protection by PML NBs favors SUMO2-dependent ubiquitination

To further establish the role of PML NBs in PML-controlled sumoylation, we investigated whether the latter depends on the ability of partner proteins recruitment which requires a specific sumoylation site in PML^6,8^. We thus generated a E167R *Pml* knock-in mouse mutated on the sumoylation consensus site and derived *Pml^E167R/-^*;His6-HA-SUMO1 mice. Critically, global hyper-sumoylation by SUMO1 and SUMO2/3 upon arsenic/pI:C co-treatment was abolished in those mice defective for partner recruitment (Fig. 2a (right part), control PML profile Supplementary Fig. 2a). Collectively, all these findings support the conclusion that PML NBs drives basal and stress-induced sumoylation of protein partners, primarily by SUMO2/3.

**Figure 2.**
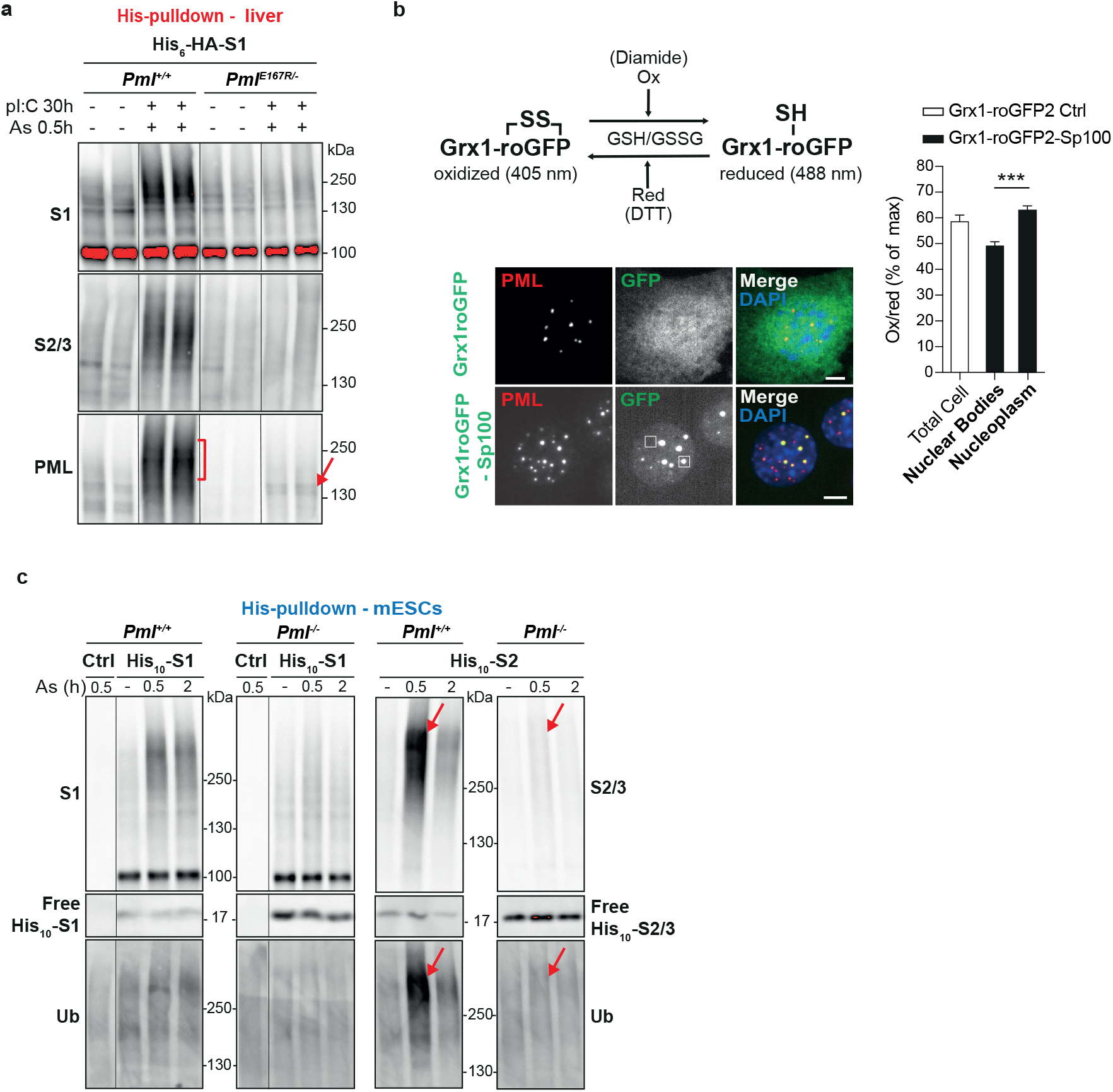
Oxidation-shielding PML NBs are required for SUMO2/ubiquitin conjugation in vivo. **(a)** Arsenic-increased sumoylation is lost in *Pml*^E167R^ knock in mice with PML NBs defective for partner recruitment, PD from the indicated mice, as in (1f). Representative experiment with 2 mice per condition, n=6. Control with PML profile from total extract in Supplementary Fig. 2a. **(b)** Reduced vs oxidized Grx1-roGFP2 sensors (top), representative images of the indicated constructs (green) in MEFs (bottom), PML (red), and regions of interest (ROI) indicative for the quantifications, scale: 5μm. Mean ratios of oxidized-405nm / reduced-488nm sensors, as percentage of max and corrected with min capacities of the sensor (right). n=3, 10 nuclei each +/− s.e.m., ***p<0.001, two-sided paired Student’s t-test. **(c)** His-PD of His_10_-S1 or His_10_-S2 conjugates from mESCs exposed to arsenic, revealing a rapid ubiquitination of the SUMO2 conjugates only, after the PML-dependent boost. The levels of the various free His_10_-SUMOs in mESCs are indicated.

In cell lines and *in vitro*, SUMO enzymes may be inactivated by ROS, which oxidize their active-site cysteines^23^. Arsenic-driven PML NB formation concentrates the SUMO-E2 UBC9 within NBs (Supplementary Fig. 2b)^8^. Since PML is a cysteine-rich oxidation-prone protein^7^, we wondered whether PML NBs might constitute a local protective environment for UBC9. We thus explored the redox status of PML NBs using the glutathione Grx1-roGFP2 sensor^28^ fused to the NB-associated SP100A protein (Fig. 2b), in which maximum and minimum oxidation were quantified upon Diamide and DTT exposure. Oxidized versus reduced Grx1-roGFP2-SP100A ratios revealed that PML NBs constitute a more reducing compartment than the rest of the nucleoplasm. Thus, the cysteine-rich shell of PML polymers may not only concentrate and immobilize UBC9 and its targets, but also shield UBC9 enzymatic activity from cellular ROS to favor processive SUMO2-conjugation.

Given that hyper-sumoylation may drive poly-ubiquitination and degradation, we assessed the fate of PML-driven hyper-sumoylated proteins upon arsenic stress *in vivo*. We found that sumoylated proteins decreased rapidly after their initial boost in *Pml^+/+^* mESCs and liver (Supplementary Fig. 2c-d). This decrease was accompanied by a PML-dependent wave of dual SUMO2-specific/ubiquitin conjugation and accumulation of poly-ubiquitinated proteins in PML NBs (Fig. 2c and Supplementary Fig. 2d-f). *In vivo* pre-treatment with bortezomib, a proteasome inhibitor, stabilized SUMO2/3 conjugates upon arsenic exposure in *Pml-*proficient livers or APL (Supplementary Fig. 2g). Collectively, these observations support the idea that PML NBs enables SUMO2-dependent ubiquitination and degradation of a substantial amount of targets under oxidative stress.

### Identification of PML NB-dependent SUMO targets

To identify the SUMO conjugates regulated upon NB-(re-)assembly, we undertook large-scale purifications of sumoylated proteins in APL mice by label-free quantitative mass spectrometry (LFQ LC-MS/MS)^25^ (Fig. 3a). Three cohorts of His_10_-SUMO2- and control APL mice were injected or not with arsenic and sampled after 1h (cohort n°1), 3h (cohort n°2) or 6h (cohort n°3). We identified reliable His_10_-SUMO2-targeted proteins, with over 60% overlap between cohorts (Table 1). In untreated mice across all cohorts, PML and PML/RARA were the top SUMO conjugates (Table 1), reflecting their known efficient baseline conjugation. Most of the *in vivo* identified SUMO2-targets in APL belonged to canonical SUMO-regulated pathways and hematopoietic regulators (Table 1), including C/EBPα, whose sumoylation drives myelo-erythroid lineage choice^29^.

**Figure 3.**
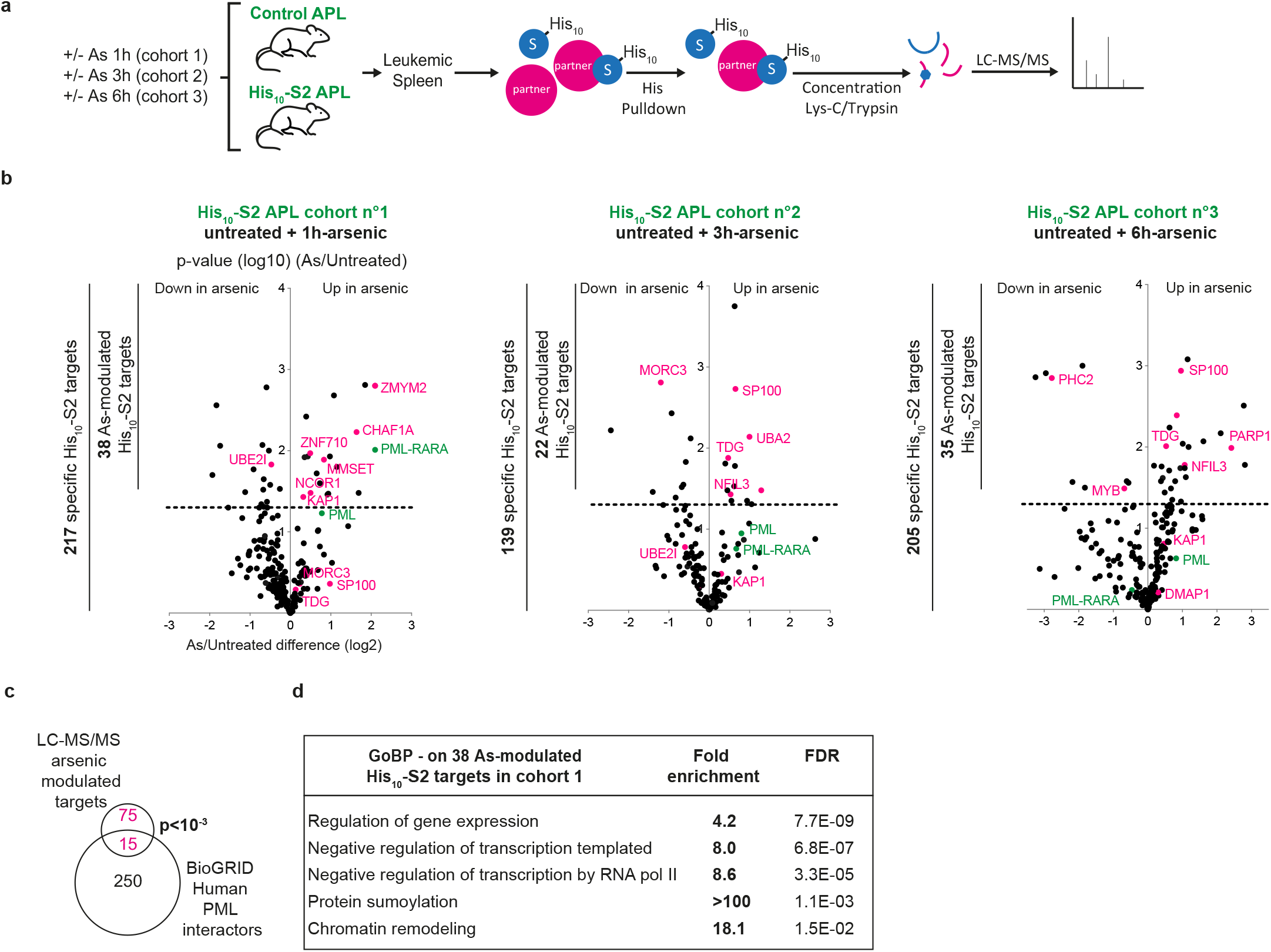
Identification of the His10-SUMO2 targets upon arsenic-induced PML NB reorganization in APL mice. Experimental setup for differential SUMO-proteome analyses upon arsenic, based on large-scale PD of His_10_-S2 conjugates from three mouse cohorts, each containing 6 Ctrl and 6 His_10_-S2 APL mice treated (n=3) or not with arsenic (n=3), for the indicated time, followed by LC-MS/MS analyses25. **(b)** Volcano plots showing proteins differentially sumoylated in arsenic-vs untreated His_10_-S2 APL mice for each cohort (label-free quantification (LFQ) fold changes, p-value with dashed line for p<0.05). Targets plotted when significantly enriched in His_10_-S2 vs Ctrl APL mice (Table 1), numbers of identified proteins in each group (left bars). **(c)** Venn diagram showing proteins common to arsenic-modulated His_10_-S2 targets identified in (b) and previously reported PML partners in30 and Biogrids. **(d)** Gene Ontology enrichment analysis (Biological Process) of 1h-arsenic-modulated targets, highlighting repressive transcriptional regulation.

We then focused on targets differentially conjugated by SUMO2 upon arsenic-driven NB reassembly (Fig. 3b). The majority of the 90 modulated targets had increased sumoylation levels (Fig. 3b, Table 2). Yet abundance of those targets was unchanged in the total APL proteome (Supplementary Fig. 3a), implying that arsenic actually enhanced their sumoylation efficiency. As expected, arsenic-modulated SUMO2 targets were enriched in known PML-interacting proteins (Fig. 3c)^30^. One hour after injection, when NBs started to re-organize, 19 targets had increased sumoylation (Fig. 3b, Tables 1-2), including PML/RARA and negative regulators of transcription or chromatin activity, such as NCORI (Fig. 3d). Strikingly, among those, we identified five proteins belonging to the same chromatin-remodeling complex involved in stem cell maintenance, comprising KAP1 (TRIM28 or TIF1beta), CHAF1a subunit of the CAF-1 histone chaperone, the MMSET histone methyl transferase (also known as WHSC1 or NSD2) and two DNA-docking KRAB-Znf proteins (ZNF710, ZNF148) (Figs. 3b, 4a and Table 2)^31,32^. Moreover, members of the KAP1 complex, not identified in this screen, are known PML NBs partners, such as the DAXX/ATRX histone chaperone or the heterochromatin protein 1 (HP1)^32–34^.

Remarkably, the repertoire of arsenic-modulated His_10_-SUMO2 targets underwent significant changes with time (Fig. 3b and Table 2), and we later identified increases in sumoylation of the SP100, TDG, MORC3 and NFIL3, known NB components (Supplementary Fig. 3b)^8^. Sumoylation of proteins linked to stem cell potential, such as PML/RARA, KAP1 or Polycomb group component (PHC2), all decreased, likely reflecting their SUMO2-triggered degradation (Fig. 3b, Tables 1-2). Several SUMO2 targets identified here -such as CCAR1, MORC3, KAP1-may contribute to the poorly understood PML control over p53 function (Fig. 3b, Table 2)^3,33,35,36^. Overall, this first unbiased exploration of stress-modulated SUMO2-conjugation *in vivo* identifies a critical epigenetic regulatory complex as a key target of the sumoylation activity of PML NBs.

### PML favors KAP1 sumoylation and repression of transposable elements in mESCs

Sumoylation is a critical activator of the KAP1 complex^31,37^ (Fig. 4a) and we thus explored the role of PML and SUMOs in KAP1-dependent epigenetic repression in mESCs. We first established that multi-SUMO2 KAP1-conjugates were enriched in *Pml^+/+^* mESCs relative to *Pml^−/−^* mESCs in basal conditions (Supplementary Fig. 4a). Moreover, arsenic sharply increased KAP1 conjugation by His_10_-SUMO2, but not by His_10_-SUMO1, in a PML-dependent manner (Fig. 4b, Supplementary Fig. 4a), in line with our above observations (Fig. 1). Similarly, in APL mice pre-treated with a proteasome inhibitor, KAP1 was also hyper-conjugated by SUMO2 upon arsenic treatment (Fig. 4c, controls in Supplementary Fig. 4b). KAP1-YFP had a nuclear diffuse localization and often overlapped with the PML NBs upon proteasome inhibition (Fig. 4d)^38^. Collectively, KAP1 behaves as a canonical PML partner, with stress-enhanced SUMO2 conjugation, a key functional modification for this master epigenetic regulator.

**Figure 4.**
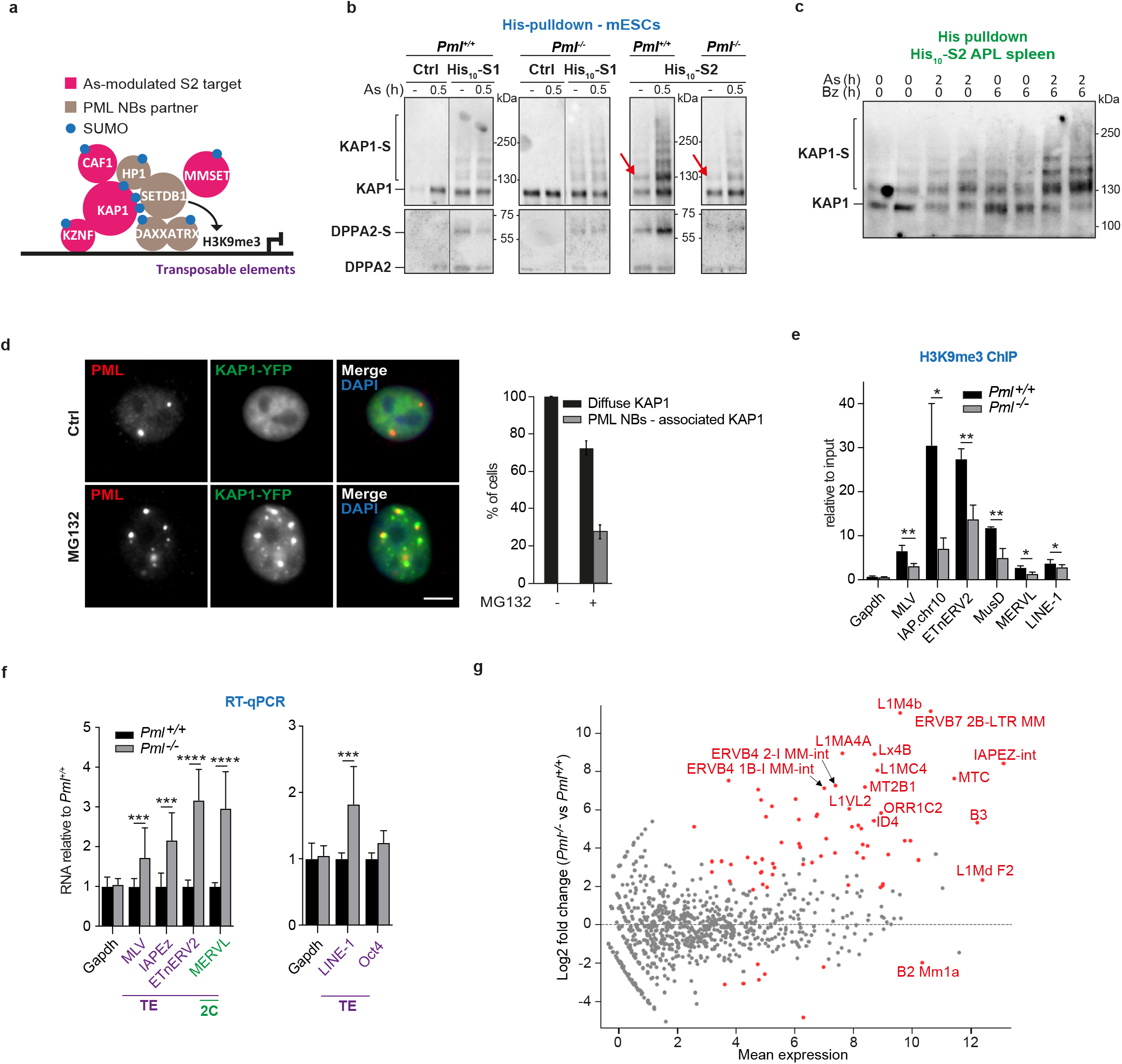
PML controls KAP1 sumoylation leading to TE de-repression in Pml−/− mESCs. **(a)** Schematic overview of the proteins in complex with KAP1. **(b)** PML-dependent arsenic-enhanced preferential S2-conjugation of KAP1 and DPPA2, PD as in Fig. 1d (see Fig. 2c for the corresponding SUMO blots). Arrows indicate the first baseline S2 adduct on KAP1. Representative data, n>3. **(c)** Arsenic-increased sumoyation of KAP1 in His10-S2 APL mice, stabilized by bortezomib pre-injection. Representative PD, see Supplementary Fig. 4b for KAP1 inputs and S2/3 conjugates. **(d)** Representative images of YFP-KAP1 (green) localization overlapping with PML NBs (red) upon MG132 in HeLa cells, scale: 5μm (left). n>3, 100 cells (right). **(e)** ChIP analysis showing decrease of the H3K9me3 mark at the promoter regions (LTRs or 5’) of the indicated TEs in *Pml*^−/−^ mESCs. Mean of n=4 replicates, relative to inputs and normalized on actin transcripts ± s.d., *p<0.05, **p<0.01 paired Student’s t-test. Representative of n=3. **(f)** Mean fold increase in the indicated TE transcripts, from n=6, normalized on actin and relative to paired values in *Pml*^+/+^ mESCs, ± s.d. ***p<0.001; ****p<0.0001, Mann-Whitney test. Representative of two CrispR/Cas9-generated *Pml*^−/−^ mESC clones. See Table 3. **(g)** MA-Plot of the RNAseq multireads analysis comparing *Pml*^−/−^ to *Pml*^+/+^ mESCs (Table 3), showing increased TE expression. Red dots highlight significant TEs with >4 fold difference.

KAP1 multi-sumoylation drives its interactions with HP1 and the SETDB1 methyltransferase, allowing trimethylation of lysine 9 of histone H3 and transcription repression in mESCs^31,32,37^ (Fig. 4a). In particular, the KAP1 complex represses endogenous retroviruses (ERVs)^39,40^, and the SUMO pathway is essential for this silencing^21,31^. First, using targeted chromatin immuno-precipitation, we observed that H3K9me3 was significantly less abundant at the 5’ LTRs of MLV, IAPEz, EtnERV2 and MusD elements of the ERV1 and ERVK families in *Pml^−/−^* versus *Pml^+/+^* mESCs (Fig. 4e). In contrast, PML-deficiency had little effect on H3K9me3 modification at the LINE-1 promoter or the MERVL LTRs (Fig. 4e), consistent with their reported indirect repression by KAP1^41,42^. Accordingly, quantification of ERV and LINE-1 transcripts by RT-qPCR demonstrated their derepression in *Pml^−/−^* mESCs (Fig. 4f). Then, in RNA-sequencing (RNA-seq) analysis for Transposable Element (TE) multi-mapped reads^43^, we found 73 repeated elements miss-regulated more than 4-fold upon PML deficiency, 68 of which were upregulated (Fig. 4g, Table 3), mostly KAP1-repressed ERVs and LINE-1. Altogether, these observations imply that PML contributes to silence TEs in mESCs, most likely through its regulation of KAP1 sumoylation and activity.

### PML controls the transition of ESCs to 2-cell like state

Many TEs are transiently expressed in totipotent 2-cell (2C) embryos, but are then repressed in pluripotent blastocysts (from which ESCs are derived). A fraction of mESCs oscillates in and out of the 2-cell like (2CL) state, recapitulating some aspects of zygotic genome activation (ZGA), a process inhibited by the KAP1 complex^44–46^. When we compared transcriptomes of *Pml^−/−^* and *Pml^+/+^* mESCs, we found highly significant enrichment in the gene set encompassing both 2C- and 2CL-restricted transcripts^44^ (Fig. 5a). Moreover, the top upregulated genes in *Pml^−/−^* mESCs include key 2CL markers such *Zscan4*, *EiF1a* and *EiF1a-like* gene families (*Zscan4b-e, Gm2016, Gm5662, Gm4340*), as well as *Piwil2*, a member of the piRNA pathway involved in TE defense^44^ (Fig. 5b, Supplementary Fig. 4c, Table 3). Expression of the *Dux* master activator, which is another hallmark of 2C embryos or 2CL mESCs^45,46^, was also upregulated in the absence of *Pml* (Fig. 5c). In contrast, transcripts of pluripotent markers, Nanog, Kfl4 or Oct4 did not significantly change with *Pml* loss in our mESCs pool (Fig. 4f, Table 3). Thus, through KAP1 sumoylation, PML may oppose the 2CL program.

**Figure 5.**
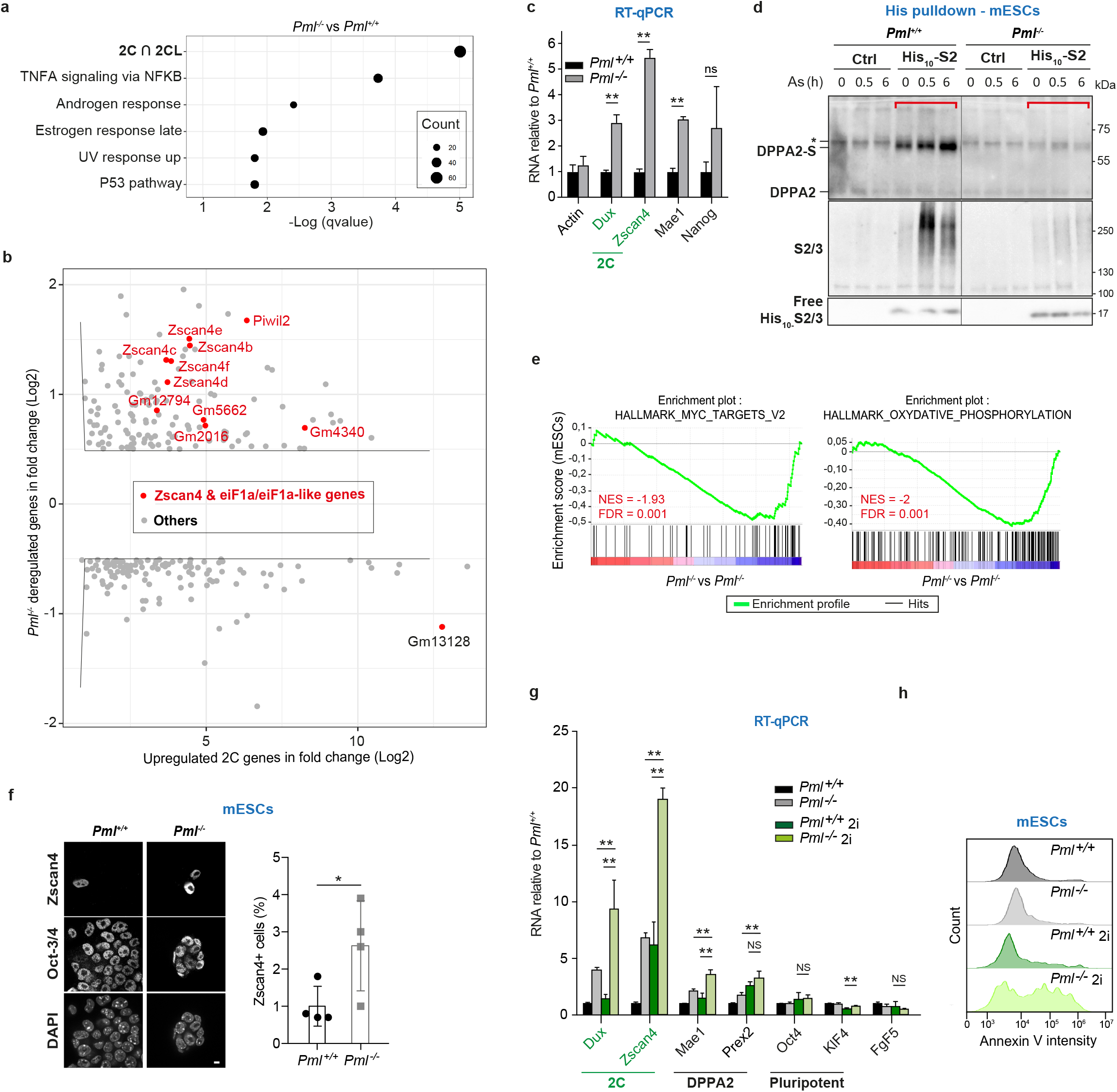
PML is required for 2CL transition and DPPA2 sumoylation in mESCs. **(a)** GSEA gene-set term enrichment analysis of the transcripts deregulated in *Pml*^−/−^ vs *Pml*^+/+^ mESCs from microarray analysis; 2C&2CL geneset: transcripts common to 2C embryo and 2CL cells from44 **(b)** Transcripts de-regulated in *Pml*^−/−^ vs *Pml*^+/+^ mESCs, related to upregulated genes in 2C embryo vs oocytes44. Transcripts with significant changes (grey) and key 2CL genes (red) are shown from microarray analysis (Table 3). **(c)** Mean fold increase of the indicated 2CL transcripts and MaeI as a direct target of DPPA2 (qRT-PCR) n=4, normalized on gapdh and relative to paired values in *Pml*^+/+^ mESCs ± s.d. **p<0.01, Mann-Whitney test. Representative of the three CrispR/ Cas9 Pml−/− clones. **(d)** PML-dependent arsenic-induced S2-conjugation of DPPA2, PD from the indicated mESCs. Inputs in Supplementary Fig. 4d. **(e)** GSEA analyses of RNAseq from *Pml*^−/−^ vs *Pml*^+/+^ mESCs (Table 3). **(f)** Representative image (right) and percentage of Zscan4+ 2CL cells (left), increased among *Pml*^−/−^ mESCs, mean ± s.d. n=4, 400-3000 nuclei; *p<0.05, Student’s t-test. **(g)** Mean fold increase of the indicated 2C transcripts (green), showing potentiation of their expression upon loss of *Pml* in ground-state mESCs cultivated in 2i media related to wildtype mESCs in FCS. n=3 **p<0.01, Mann-Whitney test. *Fgf5* is shown as a negative control of early differentiation. Representative of the three CrispR/Cas9 *Pml*^−/−^ clones. **(h)** Annexin V positive mESCs (FACS analysis), showing that loss of *Pml* induced apoptosis of ground-state mESCs.

Upstream of *Dux*, DPPA2 is another master transcription factor of ZGA, required for the establishment of 2CL state ^42,45,47^. The transcriptional activity of DPPA2 and its role to promote 2CL conversion are also blocked by its sumoylation (Yan et al., 2019). Remarkably, *MaeI* and *Prex2,* two Dux-independent targets of DPPA2^42^, were also among the top up-regulated transcripts in *Pml^−/−^* mESCs (Table 3, Fig. 5c). Critically, we demonstrated basal and arsenic-enhanced DPPA2 conjugation by His_10_-SUMO2 in *Pml^+/+^*, but not in *Pml^−/−^* mESCs (Fig. 5d, controls in Supplementary Fig. 4d). Similar to that of KAP1, His_10_-SUMO1 conjugation of DPPA2 was low, independent of *Pml* and not modulated by arsenic (Fig. 4b). Other PML-regulated SUMO2 conjugates identified in the mass spectrometry experiment, such as ZMYM2 or CHAF1a (Fig. 3b, Table 2), may also oppose 2CL transition^46,48^. *Pml^−/−^* mESCs exhibited down-regulation of MYC and oxidative phosphorylation pathways (Fig. 5e, Supplementary Fig. 4e, Table 3), another feature of 2CL cells^49,50^. Phenotypically, the number of ZSCAN4-positive and OCT4-negative 2CL cells increased with the absence of PML (Fig. 5f).

Ground-state ESCs may be obtained by triggering global DNA demethylation in 2i culture media with vitamin D^51^. In this setting, loss of *Pml* again led to a drastic derepression of 2CL genes (Fig. 5g). Interestingly, *Pml^−/−^ mESCs* stopped growing and became apoptotic when maintained in ground state, implying an essential role for PML in survival at this state (Fig. 4h and Supplementary Fig. 4f). Collectively, these data establish a key role for PML in opposing the conversion of mESCs into 2CL cells, at least in part by enforcing KAP1 and DPPA2 SUMO2-conjugation.

## Discussion

Abundance and integrity of PML NBs are challenged in a variety of physio-pathological conditions, ranging from oxidative stress to cancer or viral infections. The actual role of PML NBs was puzzling due to the functional diversity of the one-by-one identified partner proteins. Formation of PML NBs is regulated by both the abundance of PML (depending on stem cell status, interferon or P53 signaling) and its stress-responsive dynamics, as explored here with arsenic. Our findings establish the biochemical function of PML NBs *in vivo.* PML NBs drive SUMO2 conjugation at baseline in mESCs and upon stress *in vivo*, subsequently regulating partner function or stability (Fig. 6). From yeast to human, sumoylation plays a critical role in adaptive stress responses^23,52^. Distinct from earlier studies^53,54^, our differential SUMO-proteomic evaluates stress responses *in vivo*, demonstrating the key role of PML and identifying novel targets. Mechanistically, our data suggests that ROS-triggered cysteine-rich PML shells^4,7^ protect UBC9 from oxidation, and supports a model in which liquid-like condensates immobilize active enzymes and their substrates, to facilitate PTM of low abundant regulatory proteins^55,56^. In particular, PML NBs increase UBC9 processivity, enabling stress-induced global poly-SUMO2 modifications and proteasome degradation^15,16,57^ (Fig. 1–2). Other NB-associated UBC9 regulators may further enhance this activity^58–60^. In arsenic-treated APL mice, kinetic changes in the spectrum of SUMO targets suggest that the localization of PML- initially in close proximity to chromatin and later in reorganized NBs-determines partner selectivity. Given its multiple roles, the KAP1 complex could enforce leukemogenesis through PML/RARA-driven target gene silencing, but later contribute to therapy-induced PML/p53-mediated senescence of leukemia initiating cells^32,61^. Beyond APL, PML-facilitated KAP1 repressive functions may actually contribute to senescence, viral latency^62,63^ or cancer stem cell biology^12,64,65^. The proposed role of PML in chromatin status may also involved KAP1^66–68^.

**Figure 6.**
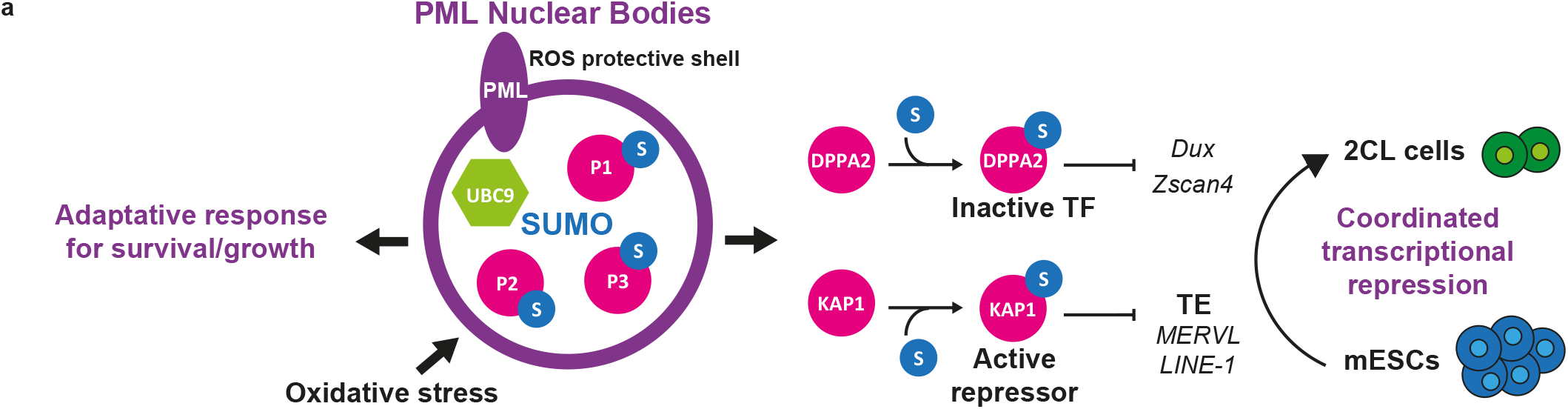
Sumoylation activity of PML NBs subsequently regulates target function, coordinating regulators of mESC fate.

Sumoylation sustains epigenetic repression in mESCs, contributing to repression of both ERVs and 2CL program^21,31,69^. The PML-dependent control of TE expression unraveled here likely explains basal activation of interferon signaling in multiple types of *Pml^−/−^* cells^70,71^ (data not shown). A previous study implicated PML in mesoderm specification at the expense to endoderm in differentiated mESCs^13^. Here we demonstrate that, by coordinating the action of various repressors and activators through the orchestration of their SUMO2 conjugation, PML NBs oppose mESC totipotent-like state and TE expression (Fig. 6). PML-dependent survival of ground-state mESCs highlights the role of PML NBs in environmental adaptation. Overall, PML NBs appear as essential regulators of mESC homeostasis, orchestrating proteostasis, cell fate and adaptive responses through sumoylation/ubiquitination control.

## Acknowledgements

This work was supported by grants from Agence Nationale pour la Recherche, ANR SUMOPiv (VLB); the ERC, PML-Therapy advanced (HDT); Fondation ARC (VLB); ANRJC (PT). We kindly thank the staff of the animal facility in Research Institute of Saint–Louis Hospital (IRSL), A.L. Maubert for her help with mouse care. We are also grateful to image core microscopy facilities of IRSL and College de France, Paris, in particular N. Setterblad for his helpful advices with incucyte analysis and A. Alberti. We warmly thank M. Tirard and N. Brose (Max Planck Institute of Experimental Medicine, Göttingen, Germany) for the sharing of His6-HA-*Sumo1* knock-in mice, and Meng-Er Huang (Institut Curie, Orsay, France) for his GRX1-roGFP2 construct. We also acknowledge P. Mayeux and V. Salnot of the Proteomic 3P5 platform in Cochin hospital, Paris, France. We kindly thank P. Lesage, A. Amara, M.H. Verhlac and D. Bourch’is for their critical readings of the manuscript, and others members of the team for helpful advices. We finally thank support services of IRSL and CIRB. The authors declare no conflict of interest.

## Author Contributions

ST, OF, MCG, performed all experiments; RGP performed quantitative MS/MS analyses, SQ analyzed RNAseq and transcriptomics data, AC and PT generated the *Pml^−/−^* mESCs, MNK the *Pml^E167R/E167R^* and *His10-HA-Sumo3* knock-in mouse, MP for *Pml^−/−^* and *Pml^E167R/E167R^* mouse crossing with His6-HA-*Sumo1* knock-in mice. OF, MCG, AV, VLB and HdT designed experiments, interpreted data and contributed to the writing of the manuscript. All authors reviewed the manuscript.

## Methods

### CRISPR/Cas9 knockout mESCs, transduction and cell culture

Mouse ESCs E14tg2a were obtained from 129/Ola blastocyst mice (P. Navarro (Pasteur, Paris) and cultured in DMEM (Gibco, #41966029) supplemented with 10% Fetal Calf Serum (FCS), 1% non-essential amino acid (Gibco, #11140050), 1% Glutamax (Gibco, #35050-038), 1% Penicillin/Streptomycin (Gibco, #10378016), 1000 units/ml of recombinant Leukemia Inhibitory Factor (LIF, Interchim, #8V6280) and 0.1% β-mercaptoethanol (Merck, #M6250) on gelatin-coated plates. Ground-state mESCs were grown in serum-free 2i medium (half NeurobasalGibco, #21103049) and DMEM/F-12 (Gibco, #11320033)) supplemented with 0.5% N2 (Gibco, #17504048) and 1% B27 supplement (Gibco, #17504044), 1% Glutamax (Gibco, #35050-038), 1% Penicillin/Streptomycin (Gibco, #10378016), 0.2 mg/ml BSA (Thermofisher, #AM2616), 0.01% Monothioglycerol (Sigma, #M6145), 10000 units/ml LIF (Merck, #8V6280), 0.1% inhibitor cocktail (Merck, #PD0325901 & CHIR99021) and 1% Vitamin C (Sigma, #A4403).

To knockout *Pml*, a single guide RNA targeting *Pml* exon 1 (Table 4) was subcloned into the pSpCas9 (BB)-2A-Puro plasmid (pX459v2, Addgene) and transfection of mESCs was performed using lipofectamine 3000 (Invitrogen, #L3000001). Puromycin-resistant colonies were expanded and three clones deficient for *Pml* were chosen after DNA sequencing and Western blot analysis.

APL cells or mESCs were transduced using retroviruses produced by Plat-E packaging cells, after transfection with MSCV-IRES-GFP constructs expressing His_10_-SUMO paralogs (Effecten reagent, Qiagen, #301425). GFP-positive cells with similar and weak fluorescent signals were sorted by flow cytometry. Transfection of pBOS-KAP1-YFP plasmid into HeLa cells was performed by using lipofectamine 2000 (Invitrogen, #11668019).

MEF, HeLa and Plat-E cells were grown in DMEM GlutaMAX (Gibco) supplemented with 10% FCS. Cells were treated with 1uM As_2_O_3_ (Sigma, #01969) or 1 uM retinoic acid (Sigma, # R2625) when requested.

### Mouse models and treatments

His_6_-HA-*Sumo1* knock-in mice deficient for *Pml* were obtained after backcrossing of 129/sv *Pml^−/−^* mice (Pier Paolo Pandolfi, USA) with C57Bl/6 His_6_-HA-SUMO1 knock-in mice (Nils Brose, Netherland) for 7 generations. Genotyping was performed by multiplex PCR using specific primers to distinguish *Pml^−/−^* (660bp) from *Pml^+/+^ (*443 bp) and tagged *Sumo1* (210bp) from endogenous *Sumo1* (163bp).

*Pml^E167R/E167R^* mice were obtained by CRISPR/Cas9 genome edition, performed on BALB/cByJ zygotes, using TAKE methods^72^. Briefly, three to four week-old BALB/cByJ females were super ovulated using CARD HyperOva (Cosmo bio, #KYD-010-EX) and human Chorionic Gonadotropin, Sigma; #CG-10) and then mated with males (8-20 weeks) to get zygotes. crRNA, TracrRNA, ssDNA and Cas9 nuclease were purchased from IDT and electroporated (NEPA21; Sonidal) to introduce *Pml* point mutation encoding E167R substitution (Table 4). Genotyping performed as described above and PCR products were sequenced (Table 4). BALB/c *Pml^E167R/E167R^* mice were then crossed with C57BL/6 *Sumo1^His6-HA/His6-HA^* mice to obtain *Pml^E167R/-^*; *Sumo1^His6-HA/wt^* heterozygote mice.

For APL mouse model, leukemic blasts of derived from h-MRP8-*PML/RARA* transgenic mice^73^ were transduced with MSCV-IRES-EGFP-His_10_-SUMO2 or -His_10_-SUMO2K/R and transplanted by intravenous (i.v.) injection into FVB mice. Serial i.v. transplantations were performed using 10^4^ GFP+ sorted APL cells from bone marrow.

7-8 week-old age and sex-matched mice were used for treatments with arsenic (5ug/g, Sigma, #202673), pI:C (20 ug/g, invivogen, #tlrl-picw), or Bortezomib (1mg/g, CliniSciences, #A10160-25) administered by intraperitoneal (i.p.) injection. Control mice were form Charles River or APL mice (untransduced blasts).Animals were handled according to the guidelines of institutional animal care committees using protocols approved by the Comité d’Ethique Experimentation Animal Paris-Nord (no. 121). Mice were maintained in a 12h light-dark cycle animal facility under specific pathogen–free conditions with free access to water and food (A03: SAFE; Institut de Recherche Saint Louis, Paris, France).

### Constructs

Coding sequences for His_10_-SUMO1, His_10_-SUMO2, or all-lysine to arginine His_10_-SUMO2 mutant (His_10_-SUMO2K/R) were PCR amplified from MSCV-HA-SUMO1 or pLV-His_10_SUMO2-Q87R or pVL-His_10-_SUMO2-allKR-Q87R constructs^8,25^ and cloned into MSCV-IRES-EGFP retroviral vector. His_10-_encoding adaptor was cloned into MSCV-SUMO1-IRES-GFP for His_10_-SUMO1construct (Table 4). For MSCV-GRX1-roGFP2-SP100 construct, pGRX1-roGFP2 plasmid was used to insert GRX1-roGFP (Meng-Er Huang, Curie Orsay, France) into pMSCV-SP100 construct^8^. For pBOS-KAP1-YFP plasmid, KAP1 was PCR amplified from pEGFP-KAP1 (Addgene, #45568) and inserted in place of PMLIII into pBOS-PMLIII-YFP plasmid.

### Incucyte cell proliferation assay

To monitor cell growth upon arsenic, 5,000 mESCs were seeded in triplicates on 96-well plates and analyzed with the IncuCyte live-cell imaging system (Essen). Four images per well were acquired with a 10X objective every 2h for a period of 2-5 days. Sequential images were then analyzed using IncuCyte software.

### Immunofluorescence and Glutathione redox potential assay

Cells were fixed in PFA 3.7% (Sigma, #HT5011) for 15 min and permeabilized in PBS 0.2% Triton X-100 for 15 min. For frozen mouse liver, 10um sections were fixed for 15 min in PFA 3.7% and permeabilized in PBS 0.2% Triton-X100 for 15 min at RT. Incubation with primary or secondary antibodies was performed in PBS 0.2 % Triton X-100 and 1% BSA for 2h before staining with DAPI for 3 min. Proximity ligation assay was performed as previously described^8^. Image acquisitions were done by confocal microscopy (Spinning disk (CSU-WI, Yokogawa) and LSM 880 (Zeiss)) or by using an Axiovert-200 inverted-fluorescence microscope (Zeiss). Images were analyzed with FIJI software.

MEFs stably expressing GRX1-roGFP2 or GRX1-roGFP2-SP100 redox sensors were seeded in glass-bottom μ-dish (Biovalley). Measurement of the glutathione redox potential was performed by confocal microscopy analysis (Spinning disk (X1/TIRF) using 405-nm and 488-nm lasers. Ratios between quantified oxidized-405nm and reduced-488nm forms of GRX1-roGFP2-SP100 were calculated within regions of interest or the whole cell. Normalization was performed using 100 uM diamide (Sigma, #D3648)) and 5 mM DTT (Sigma, #D0632) treatments to determine the maximal and minimal oxidation capacities of the GRX1-roGFP2 sensor.

### Western blot and His Pulldown

Whole cell extracts from bone marrow or spleen samples and other cell lines were lysed in 2X Laemmli buffer (Sigma,) after PBS washes. Frozen mouse liver were first homogenized using TissueLyser II (Qiagen) and then lysed in 1X Laemmli buffer, SDS-PAGE were 4-12% or 4-20% NuPAGE Bis-Tris gels (Life Technology).

For His_6_-HA-SUMO1 purification, frozen livers were homogenized as above and lysed with 6M guanidine-HCl, 100 mM Na_2_HPO_4_/NaH_2_PO_4_ pH 8.0, 10mM imidazole pH 8.0 and boiled at 95°C for 5 min. After sonication, the protein concentration was determined using ™BCA Protein Assay kit (Thermo Scientific, #23225). Lysates were equalized and His_6_-HA-SUMO1-conjugates were enriched on nickel-nitrilotriacetic acid (NiNTA) agarose beads (Qiagen, #L30210) as described in^15^. For dual purifications, the samples were then diluted with RIPA buffer, 0.4% NaDoc, 1% NP 40, 1% Triton X-100, 5mM EDTA, pH 7.5, 20mM NEM (Sigma, #E3876), PIC (Proteases Inhibitor Cocktail, Roche, #11836170001). Anti-HA Immunoprecipication was additionally performed when indicated. Samples were incubated at 4°C ON with anti-HA Affinity Matrix (Sigma, #11815016001), SUMO targets were eluted with HA peptide (Sigma, #11666975001). Pulldown of His_10_-SUMO conjugates from APL cells or mESCs at small scale was performed as above with wash buffers containing 50mM Imidazole. His_6_-MBP recombinant protein was added to the samples as an internal control of the pulldown step efficacy, when possible.

For large-scale purification of His_10_-SUMO2 conjugates, leukemic mouse spleens (with >70% of GFP+ leukemic cells) were dissociated in culture medium and washed in PBS. His_10_-SUMO2 conjugates were purified as described in^25,74^. Briefly, cells were either lysed with Laemmli buffer as inputs or stored as dry frozen pellets. Guanidine lysis buffer (above) was added to frozen pellets for sonication and protein concentration was assessed using BCA Protein Assay Reagent (Thermo Scientific, v23225). Lysates were equalized for protein concentration and incubated with NiNTA agarose beads (Qiagen, #L30210) for O/N. Beads were washed using buffer 1 with 5 mM b-mercaptoethanol and 0.1% de Triton X-100), buffer 2 (8 M urea, 100 mM Na2HPO4/NaH2PO4 pH 8.0, 10 mM Tris pH 8.0, 10 mM imidazole pH 8.0, 5 mM de b-mercaptoethanol and 0.1% Triton X-100), buffer 3 (8 M urea, 100 mM Na2HPO4/NaH2PO4 pH 6.3, 10 mM Tris pH 6.3, 10 mM imidazole pH 7.0 and 5 mM de b-mercaptoethanol) and with buffer 3 without imidazole. Samples were eluted in 7 M urea, 100 mM Na2HPO4/NaH2PO4, 10 mM Tris pH 7.0 and 500 mM imidazole pH 7.0. Eluates were passed through pre-washed 0.45 um filter columns (Millipore) to remove any residual beads and subsequently concentrated on pre-washed 100kDa cut off filters (Sartorius). Sample volume were equalized to 50 uL (5% of the samples were used as pulldown control for Westen blots analysis) and digested by LysC (Wako, #129-02541) and Trypsin (Promega, # V5111), and acidified using 2% TCA (Sigma, # T1647). Peptide samples were load on C18 StageTips and dried using vacuum.

### Mass spectrometry data acquisition

Both His_10_-SUMO conjugates were processed by mass spectrometry analyses in Alfred Vertegaal laboratory (LUMC, Netherland). All the experiments were performed on an EASY-nLC 1000 system (Proxeon, Odense, Denmark) connected to a Q-Exactive Orbitrap (Thermo Fisher Scientific, Germany) through a nano-electrospray ion source as previously described^17^. The Q-Exactive was coupled to a 15 cm analytical column with an inner-diameter of 75 μm, in-house packed with 1.9 μm C18-AQ beads (Reprospher-DE, Pur, Dr. Manish, Ammerbuch-Entringen, Germany). The gradient length was 120 minutes from 2% to 95% acetonitrile in 0.1% formic acid at a flow rate of 200 nL/minute. For the samples from cohort 2 and 3, two technical repeats were performed operating the mass spectrometer was operated in data-dependent acquisition mode with a top 5 method. Full-scan MS spectra were acquired at a target value of 3 × 10^6^ and a resolution of 70,000, and the Higher-Collisional Dissociation (HCD) tandem mass spectra (MS/MS) were recorded at a target value of 1 × 10^5^ and with a resolution of 17,500 with a normalized collision energy (NCE) of 25%. The maximum MS1 and MS2 injection times were 20 ms and 250 ms, respectively. The precursor ion masses of scanned ions were dynamically excluded (DE) from MS/MS analysis for 20 sec. Ions with charge 1, and greater than 6 were excluded from triggering MS2 analysis. For the cohort 1, the mass spectrometer was operated in data-dependent acquisition mode with a top 7 method, two independent technical repeats were performed. Full-scan MS spectra were acquired at a target value of 3 × 10^6^ and a resolution of 70,000, and the Higher-Collisional Dissociation (HCD) tandem mass spectra (MS/MS) were recorded at a target value of 1 × 10^5^ and with a resolution of 35,000 with a normalized collision energy (NCE) of 25%. The maximum MS1 and MS2 injection times were 20 ms and 120 ms, respectively. The precursor ion masses of scanned ions were dynamically excluded (DE) from MS/MS analysis for 60 sec. Ions with charge 1, and greater than 6 were excluded from triggering MS2 analysis.

### Mass spectrometry data analysis

All RAW data were analysed using MaxQuant (version 1.5.3.30) according to^75^. We performed the search against an *in silico* digested UniProt reference proteome for Mus musculus (24 March 2016) and the human PML/RARA protein.

Database searches were performed with Trypsin/P, allowing four missed cleavages. Oxidation (M) and Acetyl (Protein N-term) were allowed as variable modifications with a maximum number of 5. Match between runs was performed with 0.7 min match time window and 20 min alignment time window. The maximum peptide mass was set to 5000. Label Free Quantification was performed using the MaxLFQ approach, not allowing Fast LFQ ^76^. Instrument type was set to Orbitrap.

Protein lists generated by MaxQuant were further analyzed by Perseus (version 1.5.5.3) ^77^. Proteins identified as common contaminants were filtered out, and then all the LFQ intensities were log2 transformed. Different biological repeats of the experiment were grouped and only protein groups identified in all biological replicates in at least one group were included for further analysis. Missing values were imputed using Perseus software by normally distributed values with a 1.8 downshift (log2) and a randomized 0.3 width (log2) considering whole matrix values.

Proteins were considered to be SUMO targets when difference between His_10_-SUMO2 APL samples and their respective (Ctrl) APL control samples were statistically significant for p<0.05 (t-test) and bigger than +0.7 (log2). Thus, total SUMO targets identified in each cohort were obtained by adding specific targets found in His_10_-S2 APL *vs* Ctrl APL mice and arsenic-treated His_10_-S2 APL *vs* arsenic-treated Ctrl APL mice.

The mass spectrometry proteomics data have been deposited to the ProteomeXchange Consortium via the PRIDE^78^ partner repository with the dataset identifier PXD019609. For reviewing purposes, the following credentials can be used: **Username:** **Password:** zwrelDgt

### Chromatin immunoprecipitation

ChIP was performed with the iDeal ChIP-qPCR kit (Diagenode, #C01010180) according to the recommendation of the manufacturer. Briefly, mESC cells were crosslinked in 1% formaldehyde (Euromedex, #EM-15686) for 15 min at room temperature and quenched with 0,12 M glycine provided in the kit. The extracted chromatin was sonicated with a Bioruptor Pico (Diagenode) and immunoprecipitation was performed using an anti-H3K9me3 (Abcam, #ab8898) and a rabbit IgG-isotype (Diagenod, #C15410206) control antibody. Eluted DNA was quantified by real-time PCR on a Roche LightCycler by using FastStart Universal SYBR Green Master (Roche, #4913850001) with the set of primers listed in Supplementary Table 4. Data were normalized with respect to percentage of input and correspond to the mean S.D. from at least three replicates.

### RNA preparation and quantitative RT-PCR

Total RNA extraction from mESCs was performed with the RNeasy Plus Mini Kit (Qiagen, #74134) with an additional in-column DNase treatment. RNA was quantified using a Nanodrop One-One (Thermofisher) before cDNA amplification or RNA seq libaries preparation.

cDNA were prepared from 1 ug of total RNA with a Maxima First Strand cDNA Synthesis kit (Thermofisher, #1641) including an additional step of DNAse treatment before reverse transcription. Quantitative real-time PCR on cDNA was then performed as described for ChIP experiment. Expression levels were normalized to endogenous *Actin* or *Gapdh* gene as indicated.

### RNA sequencing

Quality control of purified RNA was performed using a 2100 BioAnalyzer (Agilent). Sequencing libraries were prepared with 800 ng of total RNA by using TruSeq Stranded Total RNA Ribo-Zero Gold Prep kit (Illumina, #20020598) allowing depletion of cytoplasmic and mitochondrial rRNA. Briefly, after reduction of rRNA with target-specific oligos combined with Ribo-Zero rRNA removal beads, purifed RNA was then fragmented and sequentially reverse-transcribed with random primers into double-stranded cDNA fragments. After adapter ligation, cDNA fragments were enriched by PCR to obtain barcoded libraries size-selected with AMPureXP beads (Beckman Coulter, #A63881). Quantification and quality control of each library was assessed by using a 2100 Bioanalyzer. Final libraries were then normalized and pooled in equal molar concentration for 75bp single-end sequencing with an Illumina NextSeq 550.

### RNA-SEQ mapping

The sequencing reads raw data (FASTQ) were submitted to quality check and were trimmed for Illumina adapters sequences and low-quality bases using Trim Galore^79^ (version 0.4.5 with default parameters). Trimmed read pairs (>35bp) were mapped to mouse reference genome (mm10) using STAR ^80^ aligner (version 2.5.2b) with maximum multiple alignments of no more than 100, using the variables *-winAnchorMultimapNmax 100* and *– outFilterMultimapNmax 100.* From BAM files, TEtranscripts ^81^ (version 2.0.3) was used to quantify both gene (uniquely aligned reads only) and transposable element transcript abundances (including both unique- and multi-aligned reads). The differential expression analysis was done by using the DESeq2 package ^82^ for modeling the counts data with a negative binomial distribution and computing adjusted *P*-values. For the comparative analyses (*Pml* ^−/−^ versus WT), only genes with both FC > 2 and FDR < 0.05 were considered as differentially expressed (Supplementary Table 3).

### Microarrays transcriptomic data analysis

Total RNA was extracted from mESCs as described for RNA sequencing analysis. Affymetrix GeneChip® Mouse Transcriptome Assay 1.0 (MTA 1.0) was used to perform gene expression analysis on 3 *Pml^−/−^* and 3 *Pml^+/+^* mESC samples. Background correction, probe set signal integration, and quantile normalization were performed through Robust Multichip Analysis (RMA) algorithm, as implemented in the “R” Affymetrix package. Genes whose abs(log2(fold-change)) between comparative group was ≥ 0.5 (with a *p* value cut-off of <0.1) were selected. In addition, false discovery rate (FDR) was applied for multiple hypotheses testing using Benjamini-Hochberg correction. Genes with a FDR-adjusted *p* value (adjusted *p* value) ≤0.05 were finally accepted. Biological gene-pathway changes were scored using the Gene Set Enrichment Analysis algorithm (GSEA; broadinstitute.org) using hallmark signature database (MSigDB). A custom specific 2C⋂2CL pathway was used based on ^44^ for transcripts restricted to both 2-cell embryo and 2CL cells.

### Statistical analysis

The number of independent experimental replications is indicated in the legends. Statistical analyses of mean and standard deviation were performed with Prism 7 (GraphPad Software) as well as Student’s t-test or Wilcoxon Mann-Whitney tests, as indicated.

## Data and code availability

The accession number for the proteomic data reported in this paper is on ProteomeXchange Consortium : PRIDE73 partner repository Identifier : PXD019609

The RNA seq (E-MTAB-10153) and Affimetrix Microarrays transcriptomic (E-MTAB-10151 data have been deposited on https://www.ebi.ac.uk/fg/annotare.

