## Supplementary Figure1 for "Unbiased *in vivo* exploration of nuclear bodies-enhanced sumoylation reveals that PML orchestrates embryonic stem cell fate"

### Tessier et al. Figure Supplementary 1

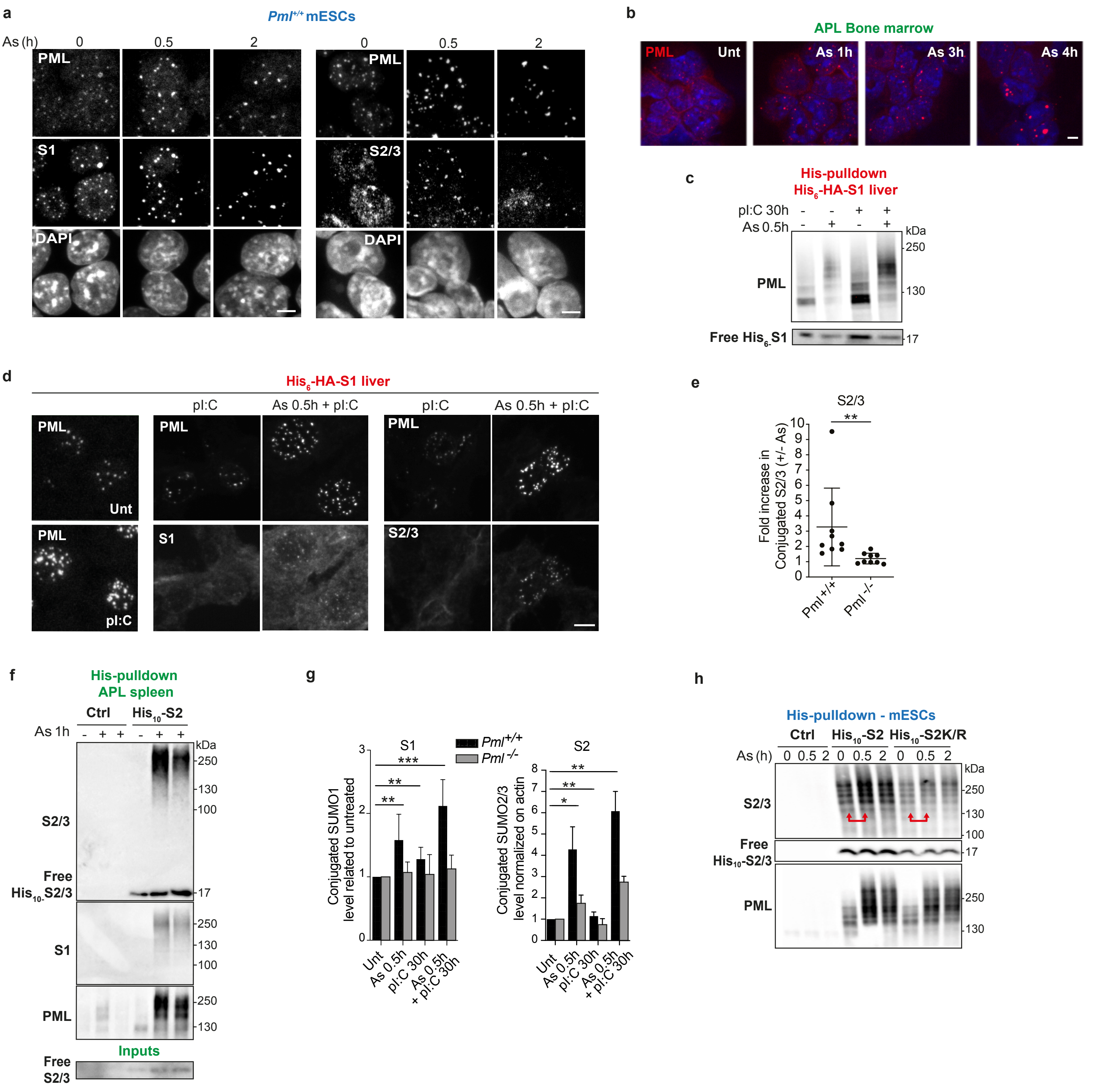


**Supplementary Figure 1.**
**(a)** Immunofluorescence analysis of mESCs showing arsenic-enhanced PML NBs and endogenous SUMO1 (lef) or SUMO2 (right) recruitment into PML NBs. **(b)** Confocal analysis of endogenous mouse PML in leukemic bone marrow from APL mice, showing fast PML NBs reorganization during arsenic kinetic. Mean intensity Z-section projections are shown. Scale bar: 5 µm. **(c)** His-pulldown from liver as indicated, showing increase in PML sumoylation and/or expression upon arsenic and/or pI:C injection, and free His6-HASUMO1. **(d)** Confocal analyses of liver cryo-sections from His6-HA-SUMO1 mice showing increase endogenous PML, SUMO1 or SUMO2/3 at PML NBs upon the indicated treatment. **(e)** Quantification of the SUMO2/3 conjugates after His-pulldown from 0.5h-arsenic-treated mESCs relative to paired untreated cells (Fig. 1d right), Mean +/- s.d. of (n=9) independent experiments. **p<0.01, ****p<0.0001 two-tailed Wilcoxon-Mann-Whitney test. **(f)** Pulldown of His10-SUMO2 conjugates from spleen of His10-SUMO2 or control APL mice, as in e, showing increased sumoylation upon arsenic-induced PML NBs reassembly. Inputs indicate unconjugated SUMO2 in APL. Representative of n=3 independent experiments with 2 mice each. **(g)** Quantification of the conjugated SUMO1 or SUMO2/3 in liver after His-pulldown and related to paired untreated mice (Fig. 1f). Data represent n >3 independent experiments, Error bars represent s.d., ** p<0.01; *** p<0.001, Fisher's exact test. **(h)** His10-SUMO2 conjugates, pulldown from mESCs stably expressing His10-SUMO2 or the all K/R mutant version, and immunoblotted with anti-SUMO2/3 and anti-PML. Red arrows underline arsenic-increased sumoylation in His10-SUMO2-expressing cells only.
