## Supplementary Figure2 for "Unbiased *in vivo* exploration of nuclear bodies-enhanced sumoylation reveals that PML orchestrates embryonic stem cell fate"

### Tessier et al. Figure Supplementary 2

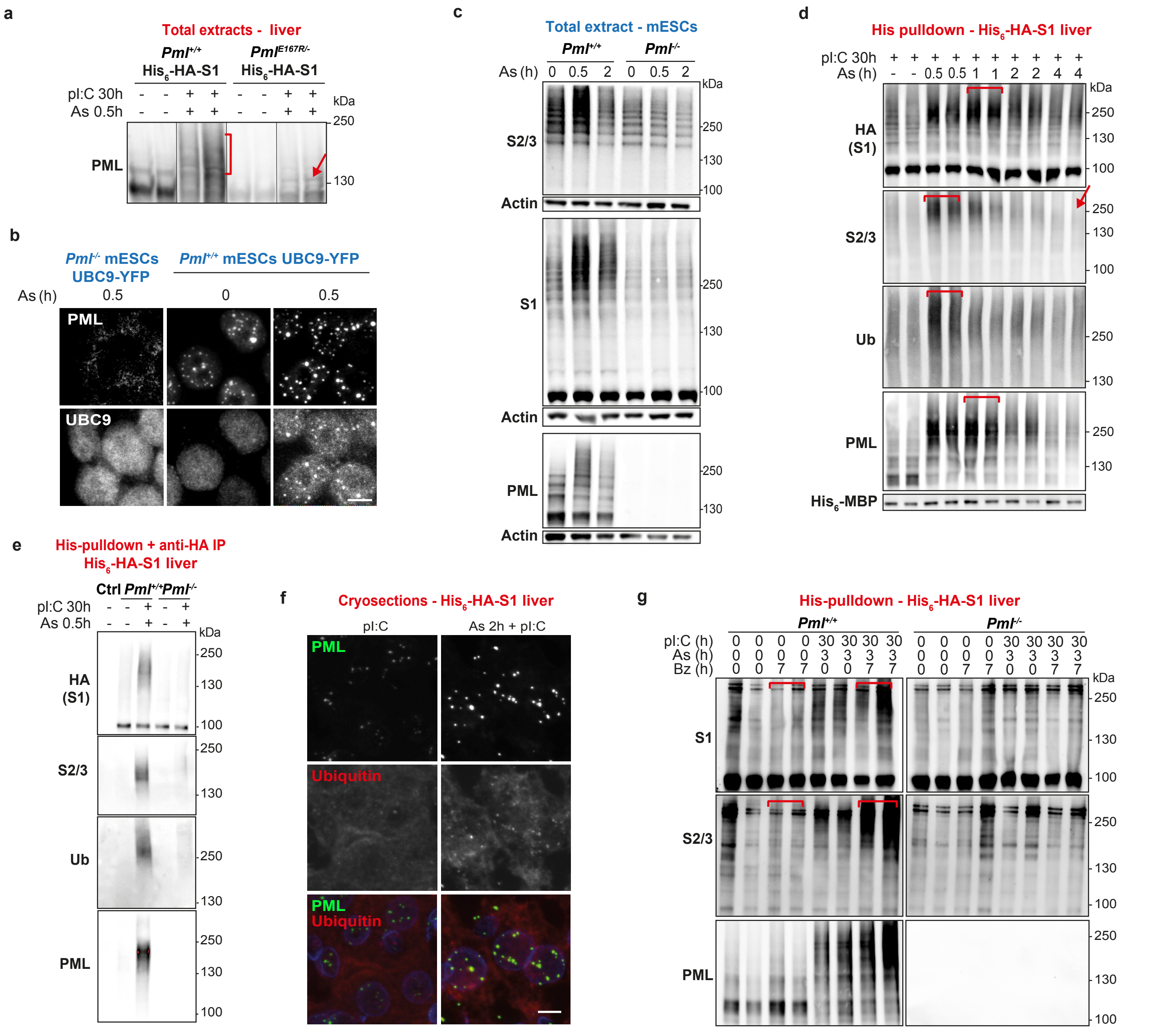


**Supplementary Figure 2.**

**(a)** Total liver extracts indicating the PML sumoylation profile in Pml+/+ or PmlE167R/- mice. n=4 independent experiments with at least 2 mice per condition. **(b)** Representative confocal images of UBC9-YFP stably expressed in mESCs showing arsenic-triggered accumulation of UBC9 into PML NBs. Z-section projections of mean intensities, scale bar: 5μm. **(c)** Total extracts of mESCs treated or not with 1μM arsenic showing endogenous SUMO2/3, SUMO1 or PML, and Actin as loading control. **(d)** Representative of His pulldown from His6-HA-SUMO1 mice liver showing wave of SUMO1 and multi-S1-SUMO2/3 or -Ubiquitin (Ub) conjugates after arsenic injection. n=2 independent experiments with at least 2 mice per condition. **(e)** His6-HA-SUMO1 conjugates from liver after consecutive His-pulldown and anti-HA IP, showing S2/3 and ubiquitin modifications in a PML-dependent manner upon arsenic injection. **(f)** Representative enrichment of ubiquitin conjugates (red) into PML NBs (green) in liver cryo-sections upon the indicated treatments. Scale bar: 5μm. **(g)** His10-SUMO2 conjugates are stabilized in APL spleen from mice pre-injected with bortezomib upon arsenic administration, two mice per condition.
