## Supplementary Figure3 for "Unbiased *in vivo* exploration of nuclear bodies-enhanced sumoylation reveals that PML orchestrates embryonic stem cell fate"

### Tessier et al. Figure Supplementary 3

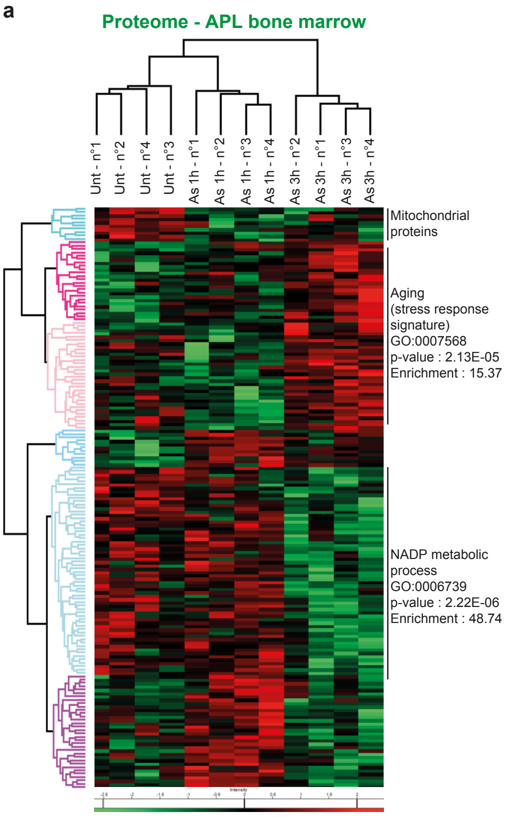


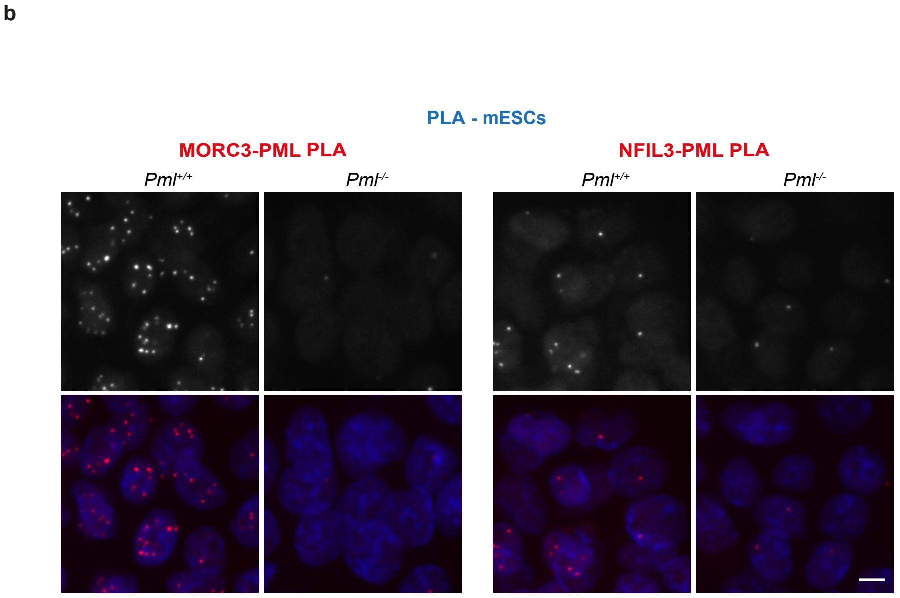


**Supplementary Figure 3.**
**(a)** Heat map analysis of total APL proteome (LFQ LC-MS/MS) upon arsenic treatment. The different clusters are indicated on the right with their associated GOBP signatures. **(b)** Proximity ligation assays showing interaction (<40nm) between endogenous PML and NFIL3 or MORC3 (red dots) in the indicated mESCs. Dapi (blue), scale bar: 5µm.
