## Supplementary Figure4 for "Unbiased *in vivo* exploration of nuclear bodies-enhanced sumoylation reveals that PML orchestrates embryonic stem cell fate"

### Tessier et al. Figure Supplementary 4

**a**

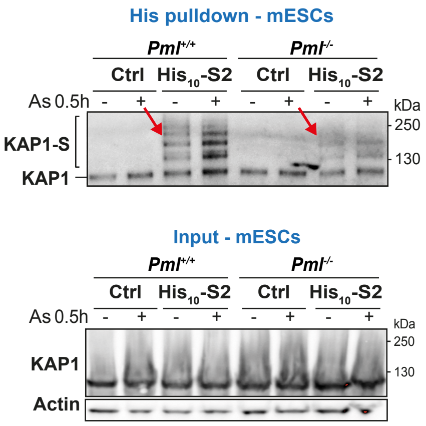


**b**

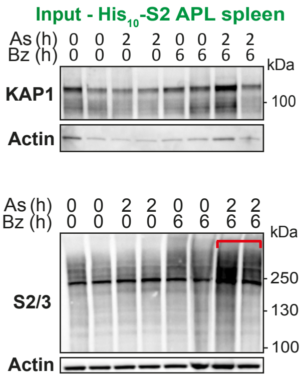


**c**

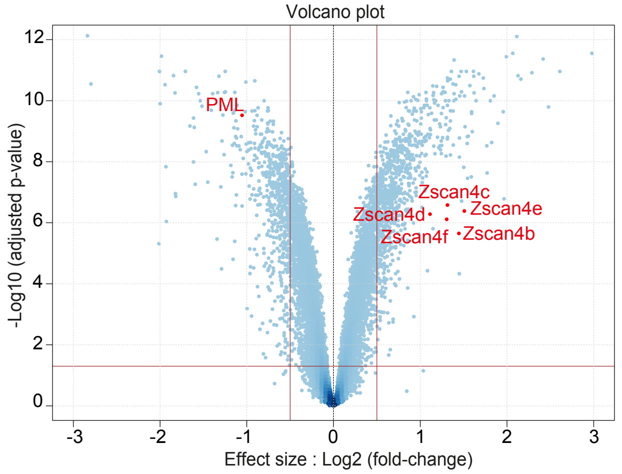


**d**

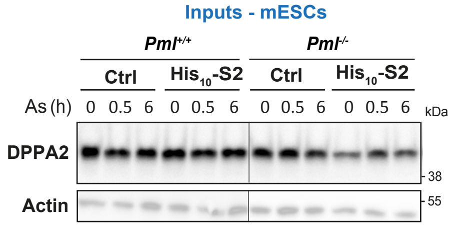


**e**

| | GS follow link to MSigDB | SIZE | ES | NES | NOM p-val | FDR q-val |
|---|---|---|---|---|---|---|
| 1 | HALLMARK_OXIDATIVE_PHOSPHORYLATION | 191 | -0.41 | -2.00 | 0.000 | 0.001 |
| 2 | HALLMARK_MYC_TARGETS_V2 | 58 | -0.48 | -1.93 | 0.000 | 0.001 |
| 3 | HALLMARK_ADIPOGENESIS | 183 | -0.35 | -1.72 | 0.000 | 0.010 |
| 4 | HALLMARK_DNA_REPAIR | 138 | -0.36 | -1.66 | 0.002 | 0.014 |
| 5 | HALLMARK_UNFOLDED_PROTEIN_RESPONSE | 109 | -0.35 | -1.58 | 0.010 | 0.027 |
| 6 | HALLMARK_MTORC1_SIGNALING | 191 | -0.32 | -1.55 | 0.000 | 0.029 |
| 7 | HALLMARK_MYC_TARGETS_V1 | 198 | -0.31 | -1.51 | 0.003 | 0.037 |

**f**

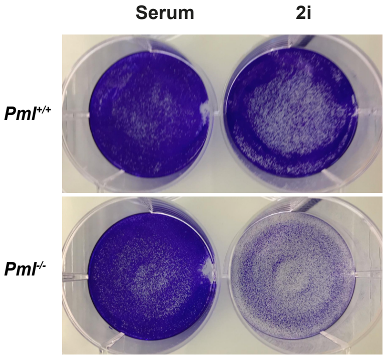


**Supplementary Figure 4.**

**(a)** Pulldown from the indicated mESCs showing PML-dependent basal and arsenic-enhanced SUMO2-conjugation of KAP1 (Top) and the corresponding inputs (Bottom). **(b)** Inputs used in Fig. 4c showing KAP1 levels (top), accumulation of SUMO conjugates in mice injected with combination of bortezomib (Bz), pI:C and arsenic, compared to Bz or pI:C/arsenic alone. Red bar underlines stabilization of sumoylation under the co-treatment compare to Bz treatment. **(c)** Volcano plot showing differentially expressed genes in Pml-/- vs Pml+/+ mESCs transcritomic analysis by affymetrix microarray (Log2 fold changes, bars indicating FC > 2, adjusted p-value < 0.01). **(d)** Inputs used in Fig. 5d showing DPPA2 levels. **(e)** GSEA analyses of RNAseq from Pml-/- vs Pml+/+ mESCs, unraveling Myc targets and Oxidative phosphorylation as the top 2 altered pathways. **(f)** Crystal violet staining of mESCs in FCS and 2i media showing that PML is essential for growth of ground-sate ESCs.
