## Supplementary Table4 for "Unbiased *in vivo* exploration of nuclear bodies-enhanced sumoylation reveals that PML orchestrates embryonic stem cell fate"

**Adaptors**

| Name | Sequence |
|---|---|
| **His10_xhol_NotI-Reverse** | CTGGTACGTAGTGGTAGTGGTAGTAGTAGTAGTAGTACGCCGG |
| **His10_xhol_NotI-Forward** | TCGAGACCATGCATCACCATCACCATCATCATCATCATCATGC |

**Primers for ChIP**

| Gene or TE name | Forward primer | Reverse primer | Reference |
|---|---|---|---|
| Actin | 5’ CCTCGATGCTGACCCTCATCC 3’ | 5’ GACACTGCCCCATTCAATGTCTC 3’ | Benabdallah NS 2019 |
| ETnERV2 | 5’ ACAAATTCAGTATGGGCATC 3’ | 5’ GGGTACTGTTAAGACCCACA 3’ | Bulut-Karslioglu 2015; Reichmann 2012 |
| ETn 5' LTR-MusD int | 5’ CCCTTCCTTCATAACTGGTGTCGCA 3’ | 5’ TAGCATCTCTCTGCCATTCTTCAGG 3’ | Thompson et al 2015 |
| Gapdh | 5’ATCCTGTAGGCCAGGTGATG 3’ | 5’AGGCTCAAGGGCTTTTAAGG3’ | Kato 2018 |
| IAPEz-chr10 | 5’ GTGCTCTGCCTTACAACTCG 3’ | 3’AAGACGCAGCAAACCAGAAT 3’ | Reichman 2011 |
| LINE-1 promoter | 5’ ACTGCGGTACATAGGGAAGC 3’ | 5’ TGTGATCCACTCACCAGAGG 3’ | Bulut-Karslioglu 2015 |
| MERVL 5' LTR-int | 5’ CTTCCATTCACAGCTGCGACTG 3’ | 5’ CTAGAACCACTCCTGGTACCAAC 3’ | Thompson 2015 |
| MLV 5' LTR-int | 5' TGGGCAGGGGTCTCCAAATCT 3’ | 5'ATAAAGCCTCTTGCTGTTTGCATC 3' | Thompson et al 2015 |

**Primers for Q-RTPCR**

| Gene or TE name | Forward primer | Reverse primer | Reference |
|---|---|---|---|
| Actin | 5’ TAGGCACCAGGGTGTGATGG 3’ | 5’ CATGGCTGGGGTGTTGAAGG 3’ | De Iaco 2017 |
| Dux | 5’ AAAGGAAGAGCATGTGCCAGC 3’ | 5’ GCAGTAAGCTGTCCTGGGAAC 3’ | De Iaco 2017 |
| ETnERV2 | 5’ ACAAATTCAGTATGGGCATC 3’ | 5’ GGGTACTGTTAAGACCCACA 3’ | Bulut-Karslioglu 2015 Reichmann et al., 2012 |
| FgF5 | 5' GCAGAAGTAGCGCGACGTTT 3' | 5' CTGGAAACTGCTATGTTCCG 3' | This study |
| Gaph | 5’ TCCATGACAACTTTGGCATTG 3’ | 5’ CAGTCTTCTGGGTGGCAGTGA 3’ | De Iaco 2017 |
| IAPEz | 5’ ACGGGAACACTTCATTACCACC 3’ | 5’ TTGAGAAGGATTCAACTGCGTG 3’ | De Iaco 2017 |
| Klf4 | 5'GAAGGGAGAAGACACTGCGT 3' | 5' GCCACTCTCCAGGTCTGTG 3' | This study |
| Lefty1 | 5' CAGCTCGATCAACCGCCAGT 3' | 5' GGCTGGCATGGCTGTGTT 3' | Kim 2014 |
| LINE-1 promoter | 5’ ACTGCGGTACATAGGGAAGC 3’ | 5’ TGTGATCCACTCACCAGAGG 3’ | Bulut-Karslioglu 2015 |
| Mae1 | 5' CGATTCCATTGCCAGGCTGC 3' | 5' AACACAGTTGCTTGGTCATGCC 3' | De Iaco 2019 |
| MERVL 5' LTR-int | 5’ ATCTCCTGGCACCTGGTATG 3’ | 5’ AGAAGAAGGCATTTGCCAGA 3’ | De Iaco 2017 |
| MLV RLTR4 MM-int | 5' AGAGGTATGGTTGGAATAAGTA 3' | 5' TAGATGGAGCCTACCAAGCTCTCAA 3' | Thompson et al 2015 |
| Nanog | 5' AAAGGATGAAGTGCAAGCGG 3' | 5' CTTCCAGATGCGTTCACCAG 3' | This study |
| Prex2 | 5' AGCATGTGAGTCTGACGGTG 3' | 5' AGGGCTTCCAGAATCTTGGC 3' | De Iaco 2019 |
| Oct4 | 5’ TCTTTCCACCAGGCCCCCGGCTC 3’ | 5’TGCGGGCGGACATGGGGAGATCC 3’ | Cossec 2018 |
| Zscan4 | 5’ AAATGCCTTATGTCTGTTCCCTATG 3’ | 5’ TGTGGTAATTCCTCAGGTGACGAT 3’ | De Iaco 2017 |

**CRISPR/Cas9**

| Target | Target sequence | ssDNA donor sequence |
|---|---|---|
| *Pml* knock-out in mESC | 5’ GCTGTGTTCATAGTCTTCGG 3’ | - |
| *Pml E167R* knock-in in mouse | 5’-CAAGCATGAGGCCCGGCCCC-3’ | 5' CAAGTGCTTCGAAGCACACCAGTGGTACCTCAAGCATagGGCCCGGCCCCTGGCTGATCTCCGCGACAATTCAGTGAGCA3' |

**Primers for His6-HA-SUMO1; *Pml*$^{+/+}$, *Pml*$^{-/-}$; *Pml*$^{E167R/-}$ mouse genotyping**

| Target | Primers for PCR | Expected size | Primer for sequencing |
|---|---|---|---|
| *Pml-/-* | 5’AAGCCATACAGGAGGAATTTCA 3’<br>5’GTGGTTGGTATTGGAGCAGAA 3’<br>5’ATCAGGATGATCTGGACGAAG3’ | *Pml*$^{+/+}$ : 443 bp<br>*Pml*$^{-/-}$ : 660bp | |
| *Pml E167R* | 5’AAGCCATACAGGAGGAATTTCA 3’<br>5’GTGGTTGGTATTGGAGCAGAA 3’ | Pml+/+ : 443 bp | 5’AGCCAAGTGCCCCAAAC3’ |
| *His6-HA-Sumo1* | 5’GGAAGTGACGCAAGACGTAGA 3’<br>5’CCCGGTACCTGGTCAGACAT3’<br>5’GAGCGTAATCTGGAACATCGTA3’ | endogenous *Sumo1* : 163bp<br>tagged *Sumo1* : 210bp | |

**Antibodies**

| Target | Source | Reference |
|---|---|---|
| Actin | Sigma | # A2066 |
| DPPA2 | Millipore | t# MAB4356 |
| HA | BioLegend | # 901501 |
| H3K9me3 | Abcam | # ab8898 |
| MBP | Sigma | # M1321 |
| MORC3 | Biotechne | # NBP1-83036 |
| E4BP4/NFIL3 | Cell signaling | # 14312 |
| PML | Millipore | # MAB3738 |
| SUMO1 | Millipore | # AB3875 |
| SUMO2/3 | Abcam | # ab81371 |
| Trim28/Kap1 | Cell signaling | # 4124 |
| Ubiquitin | Enzo | # BML-PW8810 |
| Rabbit IgG, polyclonal–isotype control | Diagenode | # C15410206 |
| Goat anti-Rabbit IgG (H+L) Cross-Adsorbed Secondary Antibody, Alexa Fluor 568 | Thermofischer | # A-11011 |
| Goat anti-Rabbit IgG (H+L) Highly Cross-Adsorbed Secondary Antibody, Alexa Fluor 488 | Thermofischer | # A-11034 |
| Alexa Fluo 488-AffiniPure Goat Anti-Mouse IgG (H+L) | Jackson immunoresearch | # 115-545-003 |
| Alexa Fluor 594-AffiniPure Goat Anti-Mouse IgG (H+L) | Jackson immunoresearch | # 115-585-003 |
| Alexa Fluor 647-AffiniPure Goat Anti-Rabbit IgG (H+L) | Jackson immunoresearch | # 111-605-003 |
| Alexa Fluor 647-AffiniPure Goat Anti-Rabbit IgG (H+L) | Jackson immunoresearch | # 115-605-003 |
